# Evolutionary and biochemical analyses reveal conservation of the Brassicaceae telomerase ribonucleoprotein complex

**DOI:** 10.1101/760785

**Authors:** Kelly Dew-Budd, Julie Cheung, Kyle Palos, Evan S. Forsythe, Mark A. Beilstein

## Abstract

The telomerase ribonucleoprotein complex (RNP) is essential for genome stability and performs this role through the addition of repetitive DNA to the ends of chromosomes. The telomerase enzyme is composed of a reverse transcriptase (TERT), which utilizes a template domain in an RNA subunit (TER) to reiteratively add telomeric DNA at the ends of chromosomes. Multiple TERs have been identified in the model plant *Arabidopsis thaliana*. Here we combine a phylogenetic and biochemical approach to understand how the telomerase RNP has evolved in Brassicaceae, the family that includes *A. thaliana*. Because of the complex phylogenetic pattern of template domain loss and alteration at the previously characterized *A. thaliana* TER loci, *TER1* and *TER2*, across the plant family Brassicaceae, we bred double mutants from plants with a template deletion at *AtTER1* and T-DNA insertion at *AtTER2*. These double mutants exhibited no telomere length deficiency, a definitive indication that neither of these loci encode a functional telomerase RNA. Moreover, we determined that the telomerase components *TERT, Dyskerin*, and the *KU* heterodimer are under strong purifying selection, consistent with the idea that the TER with which they interact is also conserved. To test this hypothesis further, we analyzed the substrate specificity of telomerase from species across Brassicaceae and determined that telomerase from close relatives bind and extend substrates in a similar manner, supporting the idea that TERs in different species are highly similar to one another and are likely encoded from an orthologous locus. Lastly, TERT proteins from across Brassicaceae were able to complement loss of function *tert* mutants *in vivo*, indicating TERTs from other species have the ability to recognize the native TER of *A. thaliana*. Finally, we immunoprecipitated the telomerase complex and identified associated RNAs via RNA-seq. Using our evolutionary data we constrained our analyses to conserved RNAs within Brassicaceae that contained a template domain. These analyses revealed a highly expressed locus whose disruption by a T-DNA resulted in a telomeric phenotype similar to the loss of other telomerase core proteins, indicating that the RNA has an important function in telomere maintenance.

## Introduction

Chromosomes in most eukaryotes are capped by tandem TG-rich DNA repeats called telomeres. The telomere repeat is remarkably conserved across deep evolutionary divergences. For example, the repeat sequence TTTAGGG present in the majority of examined plants differs by a single nucleotide from the vertebrate repeat TTAGGG [1]. Telomeric DNA repeats are recognized and bound by a suite of proteins that are essential for chromosome end protection and telomere replication [2], while telomere length is maintained by a ribonucleoprotein (RNP) complex, which minimally consists of a telomerase reverse transcriptase (TERT) and a telomerase RNA (TER). A template region within TER is complementary to approximately 1.5x the telomere repeat and is used by TERT to synthesize telomeric DNA at the chromosome terminus [3]. Although TERT and TER are sufficient to catalyze telomerase activity *in vitro, in vivo* TER serves as a scaffold for the assembly of essential accessory proteins that have a variety of functions including RNP biogenesis and recruitment of the RNP to the chromosome end [4]. Many of these accessory proteins do not interact directly, but only associate with one another via their interactions with TER [5,6]. Thus, TER is essential both as a template for telomere synthesis, and as a core scaffolding molecule in telomerase assembly.

TERT proteins from distantly related species across the eukaryotic tree are readily identifiable by homology searches based on amino acid sequence [3]. Given the essential role of TER in telomere repeat addition and binding accessory proteins, a logical expectation would be that TER would also display high levels of sequence conservation. However, TERs from different eukaryotic lineages are highly variable at the nucleotide level and appear to be entirely non-homologous, and thus independently evolved in each of the major eukaryotic lineages (i.e., animals, ciliates, fungi, and plants) [7,8]. Interestingly, most described TERs have converged on similar structural features that likely result from the shared requirement to bind both TERT and telomerase accessory proteins [6]. Two such core TER features include a pseudoknot, which is necessary for activity, and a stem loop domain (termed CR4/CR5, stem loop IV, and three way junction (TWJ) in vertebrates, ciliates, ascomycetes, respectively), which is necessary for TERT binding [9–12].

Bioinformatics approaches to recover TER have limits across major lineages of the eukaryotic crown group, but within eukaryotic clades TERs have been successfully recovered using a variety of sequence similarity based approaches. For example, in vertebrates Chen et al. (2000) recovered TER from species across the ∼450 million year radiation using sequence similarity searches and positional conservation (synteny) to identify eight conserved domains [10]. Similarly, budding yeast and its closest relatives in *Saccharomyces sensu stricto*, a radiation spanning ∼20 million years of evolution, encode TER at a syntenic locus, a fact that permitted bioinformatic detection of structurally conserved domains through signatures of covariation [13]. In the Pezizomycotina (Ascomycota), with the exception of *Saccharomyces s*.*s*., TERs display sufficient conservation to facilitate identification by BLAST using the *Neurospora crassa TER* as query. Similarly, using a combination of template sequence identification and structural motif modeling from Pezizomycotina TERs, Qi et al. (2013) recovered TERs from the more distantly related Taphrinomycotina [12]. Thus, while biochemical approaches have been required to identify TER in each major crown group lineage of eukaryotes, bioinformatics approaches have aided discovery of TER within lineages.

Until recently, the only known functional TER in plants was from *Arabidopsis thaliana*. Cifuentes-Rojas et al. (2011) found that *A. thaliana* was unusual among studied eukaryotes because it encoded two TERs: *TER1* and *TER2* [14]. *TER1* was hypothesized to serve the canonical function in providing a template for telomere addition by telomerase *in vivo. TER2* has the same template domain as *TER1*, but it is encoded at a different locus and has been hypothesized to have an alternative function in regulating telomerase activity during genome stress caused in part by double strand DNA breaks [15,16]. Both *AtTER1* and *AtTER2* were shown to assemble *in vivo* into a telomerase RNP that includes *TERT*, although the accessory proteins with which they assemble differ [14]. Importantly, the *TER2* RNP has not been observed to contribute to telomere length maintenance.

Attempts to use bioinformatics approaches to identify the TER-encoding locus or loci similar to *AtTER1* or *AtTER2* from other Brassicaceae have yielded surprising results. For example, in 15 sampled species of the family, spanning 60 million years of evolution [17], there is a single locus with sequence similarity to both *AtTER1* and *AtTER2* [18]. Sequence alignment of the recovered *AtTER1/2-like* loci from the 15 sampled species revealed changes at the template domain in four and the complete absence of a template-like domain in three. In order for the four loci with template mutations to encode for TER, a corresponding change in the telomere repeat should be observed [18]. However, genomic data from several of these species, including *Arabidopsis lyrata* and *Capsella rubella*, indicate the plant telomere repeat is conserved [19], suggesting the presence of an alternative TER locus. Although TER is known to evolve rapidly at the sequence level in other systems, the rapid rate of novel TER incorporation suggested by the identification of *AtTER1* is unprecedented. Recently, Fajkus et al. (2019) showed that *AtTER1* mutants do not exhibit telomere shortening, and thus does not act as a bona fide telomerase RNA [20].

Here we generate double mutants at *AtTER1* and *AtTER2* and verify that *AtTER2* does not play a compensatory role in telomere length maintenance in the absence of *AtTER1*. Due to the previously documented rapid evolution at the *AtTER1/2-like* locus [18], we sought to determine the evolutionary conservation of the telomerase complex using a series of comparative evolutionary and biochemical analyses on the protein constituents of telomerase. Interestingly, rather than concluding that TER and components of telomerase are evolving rapidly as the evolutionary history of *AtTER1/2* suggested, we find that the protein components of telomerase are highly conserved, as is the biochemical activity of telomerase. We performed an RNA immunoprecipitation using core protein components of the telomerase holoenzyme, followed by RNA-sequencing, and then used our evolutionary data to constrain possible template-bearing TER candidates to those that were conserved across the Brassicaceae family. This alternative approach led to the identification of the conserved plant TER discovered by Fajkus et al. (2019). Our data indicate that telomerase core components are highly conserved and our data complements the findings in Fajkus et al. (2019) in verifying that the biochemical activity of the holoenzyme is also conserved. Finally, we explore the possibility that the previously documented *AtTER-like* locus in Brassicaceae shares the *AtTER2* function of modulating telomerase activity during genotoxic stress.

## Results and Discussion

### Double *AtTER1/AtTER2* mutants lack a telomere phenotype

Beilstein et al. (2012) showed that the *TER1/2-like* locus was not conserved throughout the Brassicaceae family, or even in other species of the genus *Arabidopsis*. This led us to re-examine the conservation of *AtTER1* among the ecotypes of *A. thaliana*. Using genomic data from 1001genomes.org, we generated an alignment of the *AtTER1* locus using MEGA5 and sorted through the template domain in Geneious (Biomatters, Inc.) [21,22]. We found that *AtTER1* had a mutated template domain in 41/853 *A. thaliana* ecotypes (TCCCAAA**T** -> TCCCAAA**A**), which would preclude the use of an *AtTER1* transcript for telomere elongation in these ecotypes. Until the recently published work of Fajkus et al. (2019), previous characterization of the *AtTER1* phenotype was performed using an RNAi knock-down approach [14]; therefore, we generated a template deletion mutant at *AtTER1*.

A telomerase RNA cannot perform its templating function without an intact template domain, and thus we independently used CRISPR-Cas9 nuclease and CRISPR-Cas9-D10 nickase to create full template deletions at the *AtTER1* locus. We recovered two alleles that lacked the template domain in its entirety (Fig 1A), *Atter1 Δ18* and *Atter1 Δ22*. First generation homozygous mutants showed no reduction in telomere length as measured by telomere restriction fragment length analysis (TRF) (Fig 1B). We then propagated both *ter1* homozygous template nulls for several generations and measured telomere length in order to determine whether there was progressive loss over time. We observed no reduction in telomere length for either template-null mutant allele, regardless of generation (Fig 1B). Thus, our data definitively indicate that *AtTER1* alone does not function as the template for telomere elongation, confirming the findings in Fajkus et al. 2019.

**Fig 1.**
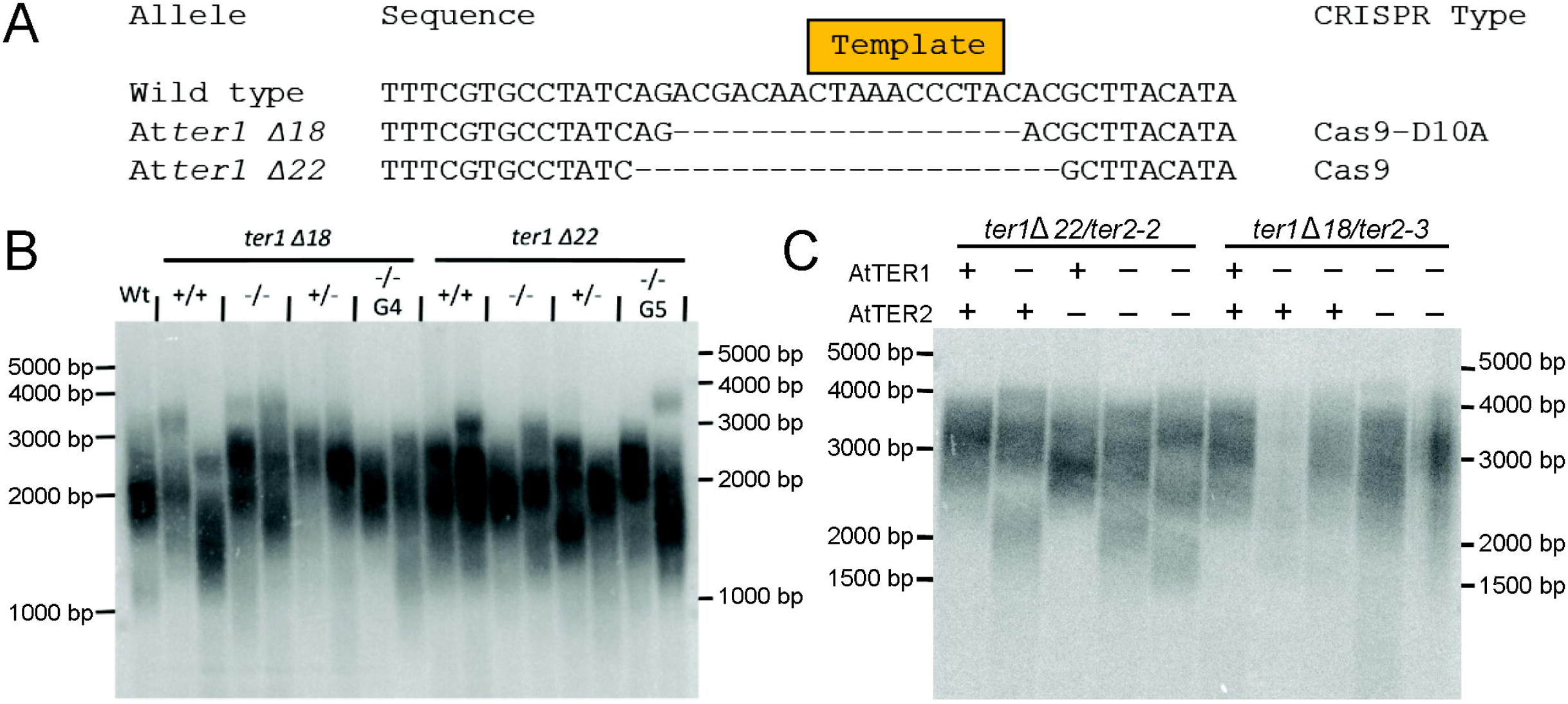
Double knockout of *TER1* and *TER2* does not decrease telomere length. (A) Two *Atter1* mutants were created using either CRISPR-Cas9 nuclease or nickase. The complete deletion of the template domain precludes the use of *AtTER1* as the template for telomere extension. (B) TRF analyses were performed on individual plants with two biological replicates in a population segregating the indicated *ter1* allele (Δ18 or Δ22), and in successive generations (G4 and G5). (C) TRF analyses were performed on two sets of double mutant (*ter1 Δ22/ter2-2* and *ter1 Δ18/ter2-3*) populations segregating the indicated mutations. A ‘+’ indicates a homozygous wild type individual for the indicated gene and a ‘-’ represents a homozygous mutant for the indicated gene.

To rule out the possibility that *AtTER2* acts in a compensatory role to elongate telomeres in the absence of *AtTER1*, we crossed both *Atter1* template null alleles (*Atter1 Δ18* and *Atter1 Δ22*), with two *Atter2* T-DNA mutants (*Atter2-2* and *Atter2-3*), to create homozygous double mutants and then measured telomere length by TRF. Neither single mutants for either gene, nor double mutants showed a decrease in telomere length (Fig 1C). Taken together, the lack of decrease in telomere length in both the single and double mutants indicates that *A. thaliana* requires neither the *AtTER1* nor the *AtTER2* locus to elongate telomeres.

### Telomerase RNP evolution is under purifying selection in Brassicaceae

Relatively high levels of conservation of the telomerase RNA within major eukaryotic lineages has allowed for the identification of distantly related TERs within those lineages. It was the unprecedented lack of conservation of the template domain at *AtTER1* within *A. thaliana* and *AtTER1/2-like* locus within other species of Brassicaceae that motivated further exploration of telomerase RNP evolution and function. Numerous genome expansions and contractions within the Brassicaceae family [23] have created opportunities for gene neofunctionalization or subfunctionalization, potentially leading to incorporation of novel subunits into the telomerase RNP in Brassicaceae, including the potential for alternative TERs [18]. If novel subunits have been incorporated into the telomerase RNP, we might expect the core subunits to undergo structural or chemical changes associated with the accommodation of novel binding partners. These changes would be expected to arise under positive selection. Alternatively, if the telomerase RNP has not undergone this type of change, we would expect the core subunits to evolve under purifying selection. With this in mind, we sought to determine if major protein components of the telomerase RNP exhibited signatures of positive selection, indicating underlying changes to the complex, or purifying selection, indicating a stable RNP. We tested for positive selection in *TERT* along several branches within the Brassicaceae. In total, we tested five branches indicated by roman numerals in Figure 2 using the branch-sites test in PAML 4.4b [24]. In each case we recovered no evidence that *TERT* evolved under elevated rates of non-synonymous to synonymous substitutions (Fig 2), suggesting that *TERT* remains under strong purifying selection.

**Fig 2.**
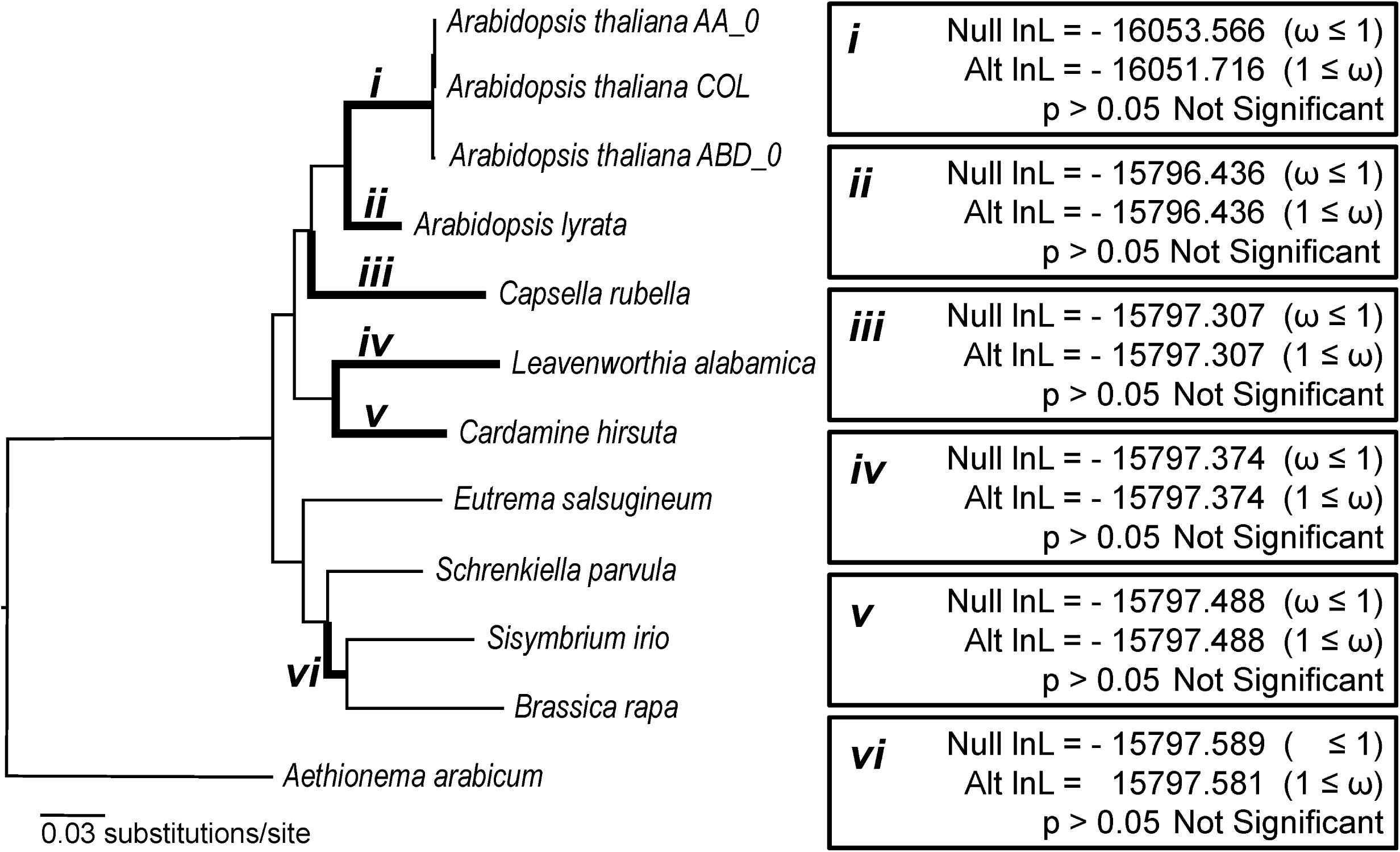
Tests for signatures of positive selection in TERT evolution in Brassicaceae. Branches throughout the Brassicaceae phylogeny, indicated by italicized roman numerals and bolded branches, were tested for positive selection, Likelihood ratio tests were performed on the bolded branches using PAML (codeml) to compare the likelihood scores of models of evolution that either include site classes with ω values ≥ 1 (Alt = Alternative model) or explicitly exclude these site classes (Null = Null model). Alternative model likelihood scores that are ≥ 1.92 better than null model likelihood scores indicate a significant signal of positive selection (p<0.05) along the specified branch (denoted as *i*-*vi* on the tree).

In addition to *TERT*, we also searched for evidence of positive selection in three other genes that encode telomerase accessory factors in *A. thaliana*. Dyskerin is a pseudouridine synthase necessary for maturation of rRNA that has also been shown to be an essential component of telomerase in *A. thaliana* and many other eukaryotes [25]. KU is a heterodimer composed of KU70 and KU80 that associates with telomerase in humans, budding yeast, and *A. thaliana* via Ku80-telomerase interactions [15,26,27]. KU plays dual roles as a key factor in double-strand DNA break repair via the non-homologous end-joining pathway, as well as telomere maintenance [28,29]. With the exception of the branches leading to *C. rubella* (iii) and *S. irio/B. rapa* (vi) for *DYSKERIN* and *A. lyrata KU70*, we found no evidence of positive selection (S1 Figure). Thus, contrary to our initial expectations, the protein components of telomerase are not responding at the molecular level to genome duplications or contractions, such as those that occurred in species like *B. rapa*. These findings support the idea that a highly conserved TER locus may be present and functional in Brassicaceae.

### An evolutionary analysis of Brassicaceae telomerase enzymology recapitulates the accepted organismal phylogeny

We next sought to take an indirect, but more fine-scale evolutionary look at how conserved TERs are in Brassicaceae, specifically in the region within and adjacent to the template domain. The ability of telomerase to bind and extend a substrate comes from TERT-substrate and TER-substrate interactions [8]. In addition, telomere synthesis requires Watson-Crick base-pairing between the 3’ end of the DNA substrate and the beginning of the template domain within TER [30]. Thus, changes within and surrounding the template domain can lead to differential alignment on a telomere substrate or premature product dissociation, particularly when these changes occur within the 3’ end of the template region [31–33]. With this rationale in mind, we set out to determine if the origin of the structurally similar Brassicaceae TERs came from one evolutionary event (i.e., an alternative TER locus with shared ancestry for all Brassicaceae) or multiple independent evolutionary events (i.e., alternative TER loci in each species that converged rapidly on a similar TER structure).

To distinguish between these two possibilities, telomerase substrate utilization was performed in ten species across Brassicaceae (Fig 3; S2 Figure). To generate a profile of substrate utilization, we used the telomere repeat amplification protocol (TRAP) on partially purified plant extracts incubated with a suite of oligos of different lengths and 3’ end composition, ranging from one to three nucleotides capable of Watson-Crick base-pairing with a TER template (Fig 3; S2 Figure) [34]. Radio-labeled products were separated on a polyacrylamide sequencing gel and compared against an oligo expected to produce the shortest telomere permutation based on previous observations (N_15_-GGG; Fig 3) [35]. Observed differences in size between products relative to this oligo were measured and recorded as a value of 0-6 using *A. thaliana* as the baseline (Fig 3, S2A Figure). A substrate utilization profile was then generated for nine other species within Brassicaceae, including several close relatives to *A. thaliana* (three biological replicates per species; summarized in Fig 3; S2B Figure). This profile was then compared against the phylogenetic tree reflecting the known relationships among these species (Fig 3).

**Fig 3.**
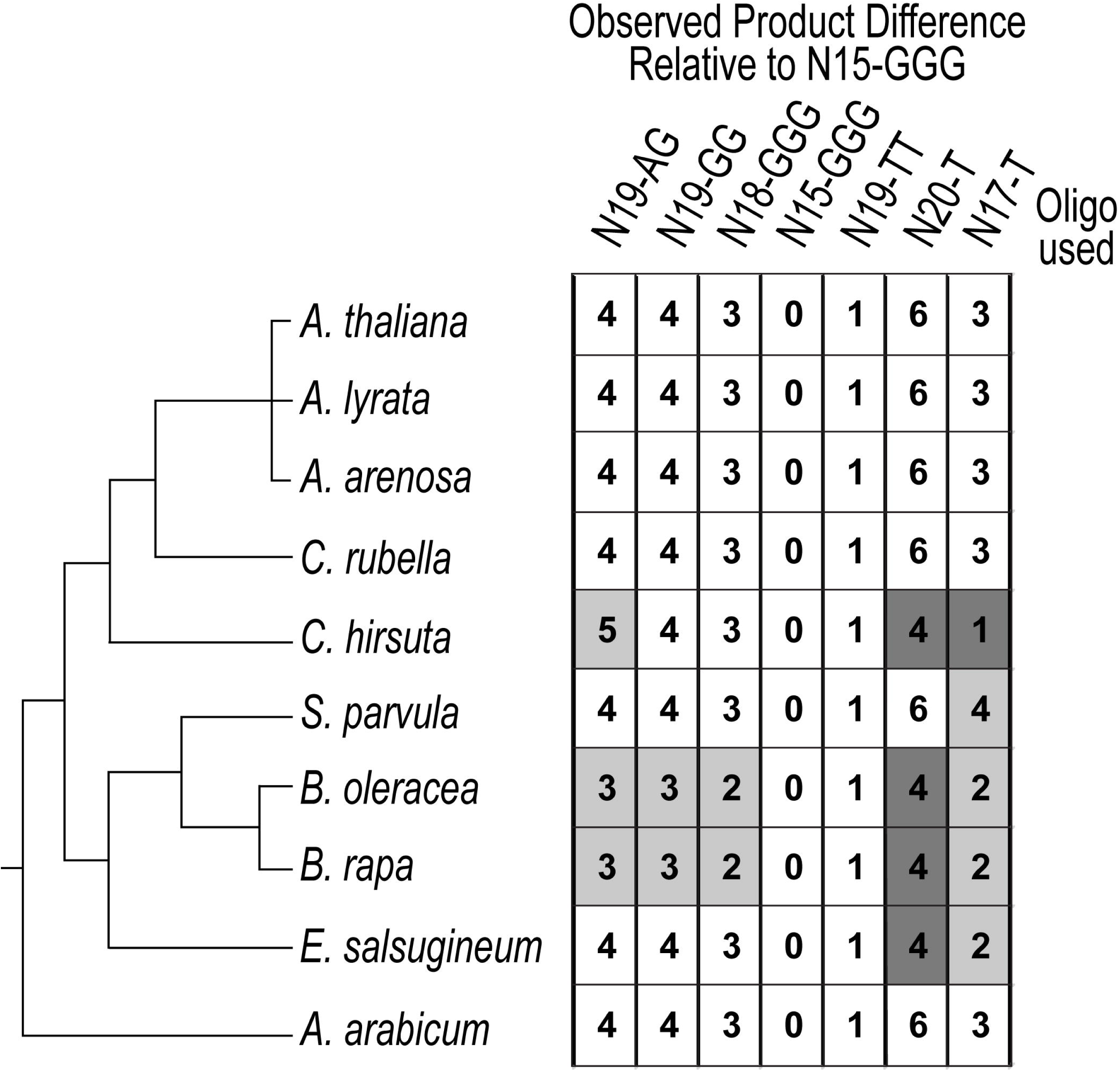
Substrate utilization by Brassicaceae telomerase RNPs closely resembles the organismal phylogeny. Substrate utilization profile for telomerase RNPs across ten Brassicaceae. Left, the accepted organismal phylogeny of species examined. Right, substrate utilization profile for all tested species. Telomerase extracts were incubated with a suite of oligos of varying length and nucleotide composition. The site of binding and number of nucleotides added prior to the first full repeat dictate the size of products produced. The lengths of each product were determined by comparing against the shortest permutation (N15-GGG). Observed product differences relative to N15-GGG were calculated for all species and oligo combinations. The intensity with which the numbered boxes are shaded corresponds to the degree to which the results differ from A. thaliana.

The substrate utilization profiles we generated for Brassicaceae telomerase closely recapitulated the evolutionary history of the family (Fig 3). The Arabidopsis clade, along with *C. rubella*, all utilized the suite of oligos in the same manner (White boxes; Fig 3). Importantly, *A. arabicum*, which represents the earliest diverging lineage in Brassicaceae [36], shares the profile of *A. thaliana*, despite 55 million years of evolution. Thus, these data support a model for shared ancestry of a TER locus in all Brassicaceae.

### Distantly related Brassicaceae TERTs assemble into a functional telomerase RNP in *A. thaliana*

To confirm our evolutionary analyses indicating that both the protein and RNA components of the telomerase RNP are likely highly conserved across Brassicaceae, we attempted to complement *A. thaliana tert* null lines with the genomic version of *TERT* from four species in the group: *Capsella rubella, Cardamine hirsuta, Eutrema salsugineum*, and *Schrenkiella parvula* [37]. The ability to complement *A. thaliana tert* null mutants with TERTs from divergent species would indicate high conservation of telomerase across Brassicaceae. In *A. thaliana*, plants can survive for approximately nine generations in the absence of TERT [38]. During this time, telomeres progressively shorten until they reach a length sufficient to elicit a DNA damage response that prohibits further cell division. We determined the degree to which the *Attert* null was complemented by each of the transgenes using a chromosome arm-specific telomere length assay known as PETRA [39,40]. In the wild type *A. thaliana* (Col-0) background, telomeres for all three arms tested were between the typical 2-4 kilobase size range (Figs 4A and 4C) [39]. As expected, telomeres in the unselected sixth generation *Attert* -/- mutants were between 0.5-2 kilobases in length (Figs 4A and 4C). The *AtTERT* transgene was able to complement the mutant background and restore telomeres back to wild type range (Figs 4A and 4C; three independent biological replicates for each construct). Telomeres in these lines were elongated further in the second generation, suggesting successful and complete complementation with this construct (Figs 4A and 4C). We observed similar or slightly better complementation with TERT from *S. parvula*, one of the most distantly related species in our study (Figs 4B and 4C) [17]. Interestingly, despite observing transcription of the *E. salsugineum, C. hirsuta*, and *C. rubella TERT* transgenes (S4 Figure), and telomerase activity in these complementation lines, we did not observe telomere elongation with these constructs (S3 Figure). However, with the exception of the 1L chromosome arm in *C. rubella*, there was no significant decrease in telomere length between the two generations we tested (S3B Figure), suggesting some degree of partial complementation.

**Fig 4.**
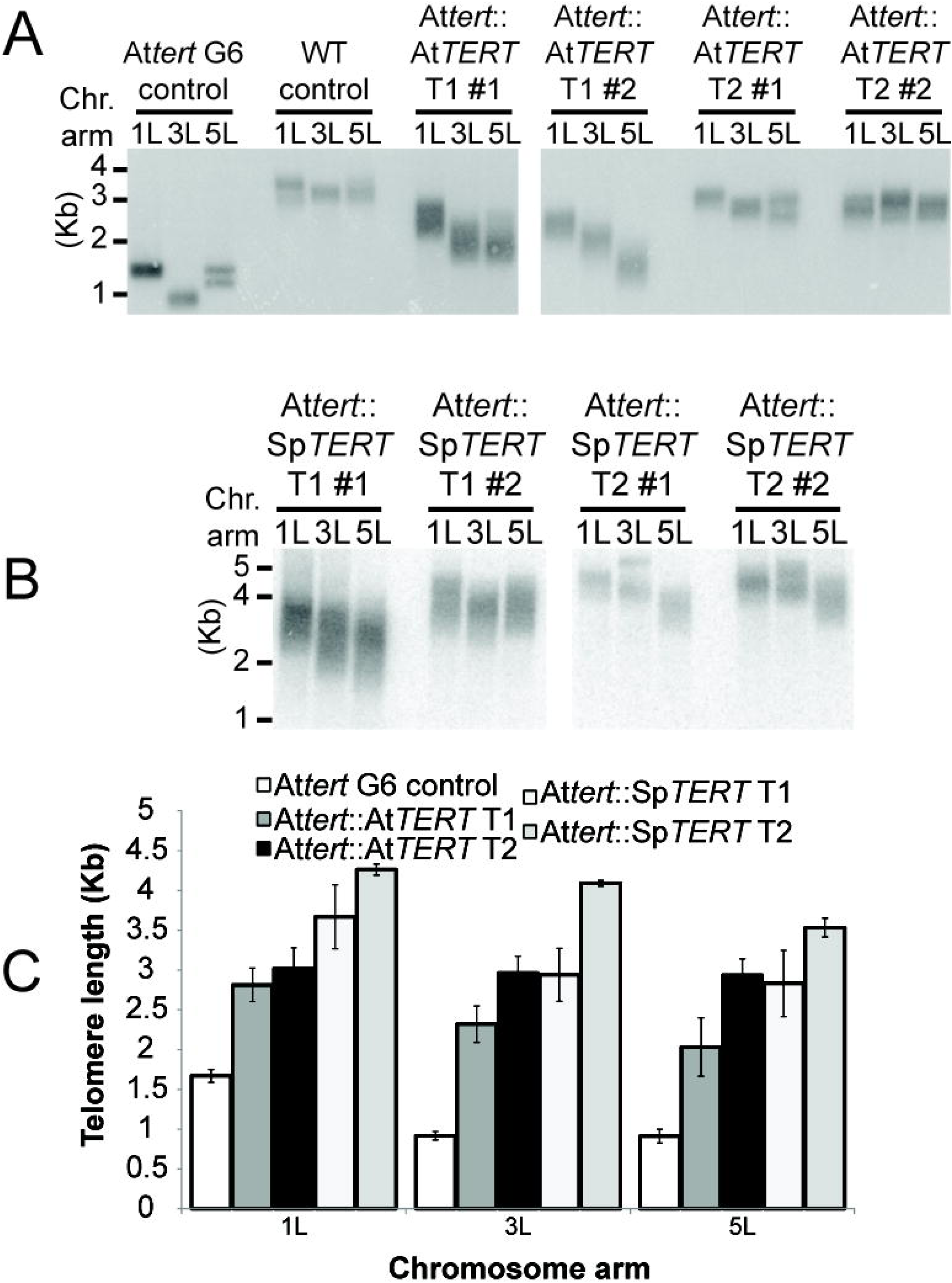
*S. parvula* TERT complements loss of function *Attert* -/- mutants. (*A*) PETRA results of the unselected, untransformed (US) control, wild type *A. thaliana* (Col.), and *Attert* -/- mutants transformed with the *AtTERT* construct. Chromosome arms are depicted by 1L, 3L, and 5L and indicate the left arm of chromosomes 1, 3, and 5, respectively. T1 and T2 indicate the first and second generation after transformation. At least three independent T1 transformants were analyzed and then propagated to the second generation. (B) PETRA results of complementation of *Attert* -/- mutants by *S. parvula* (Sp) TERT. (C) Quantification of PETRA results from (A) and (B).

These data may represent an inability on the part of some of these *TERT*s to fully reconstitute telomerase in *A. thaliana*. However, *C. rubella, C. hirsuta*, and *E. salsugineum* all have short endogenous telomeres, ranging from 1-3 Kb [19]. *S. parvula*, in contrast, has telomeres closer in length to *A. thaliana*. Thus, what appears to be partial complementation in *A. thaliana* may reflect some feature in the *C. rubella, C. hirsuta*, and *E. salsugineum* TERT protein that governs the production of shorter telomeres. The observed complementation in *A. thaliana* by TERTs from other Brassicaceae supports the hypothesis of a common TER secondary structure in all Brassicaceae.

### Identification of an alternative telomerase RNA by immunoprecipitation of the telomerase RNP followed by RNA-seq

CRISPR induced template deletion mutants of both previously described TER loci in *A. thaliana* indicated that neither is a bona fide TER. We performed a series of immunoprecipitation (IP) experiments designed to purify the telomerase RNP from floral tissue, seedling tissue, and plant cell culture, all of which express *TERT* at a high level [37]. We tracked telomerase activity from our IP and input fractions using TRAP, and extracted RNA from IP fractions with verified telomerase activity. Following cDNA synthesis we pooled and sequenced the libraries from IP experiments that used either an anti-TERT or anti-POT1A antibody. Sequencing yielded 164,505,451 total paired-end reads across the 20 IP experiments. The resulting reads were mapped to the *A. thaliana* genome (TAIR 10) and long non-coding RNAs (lncRNAs) were identified using Evolinc (Fig 5A) [41,42]. Interestingly, no reads mapped to the *AtTER1* locus. We winnowed candidate TER loci by sorting through Evolinc predicted lncRNAs to find those capable of generating the telomeric repeat and those that were conserved across Brassicaceae, in accordance with our evolutionary analyses (Figs 2-4) suggesting a highly conserved TER. From the candidate set, we identified five highly-expressed loci exhibiting conservation across the family (Fig 5B). The locus with the highest average TPM among those conserved in at least three of our representative Brassicaceae is the same locus that was identified as a telomerase RNA, named *TR*, which was shown to be conserved across land plants [20]. Moreover, we were further prompted to analyze the *TR* locus based on data produced by the Shippen Lab at Texas A&M University as part of our collaborate efforts to identify the bona fide *AtTR* (see Song et al., in revision). This locus was also previously identified as a lncRNA associated with hypoxic stress, termed *AtR8*. Hereafter we will refer to this locus as *AtTR/R8* [43].

**Fig 5.**
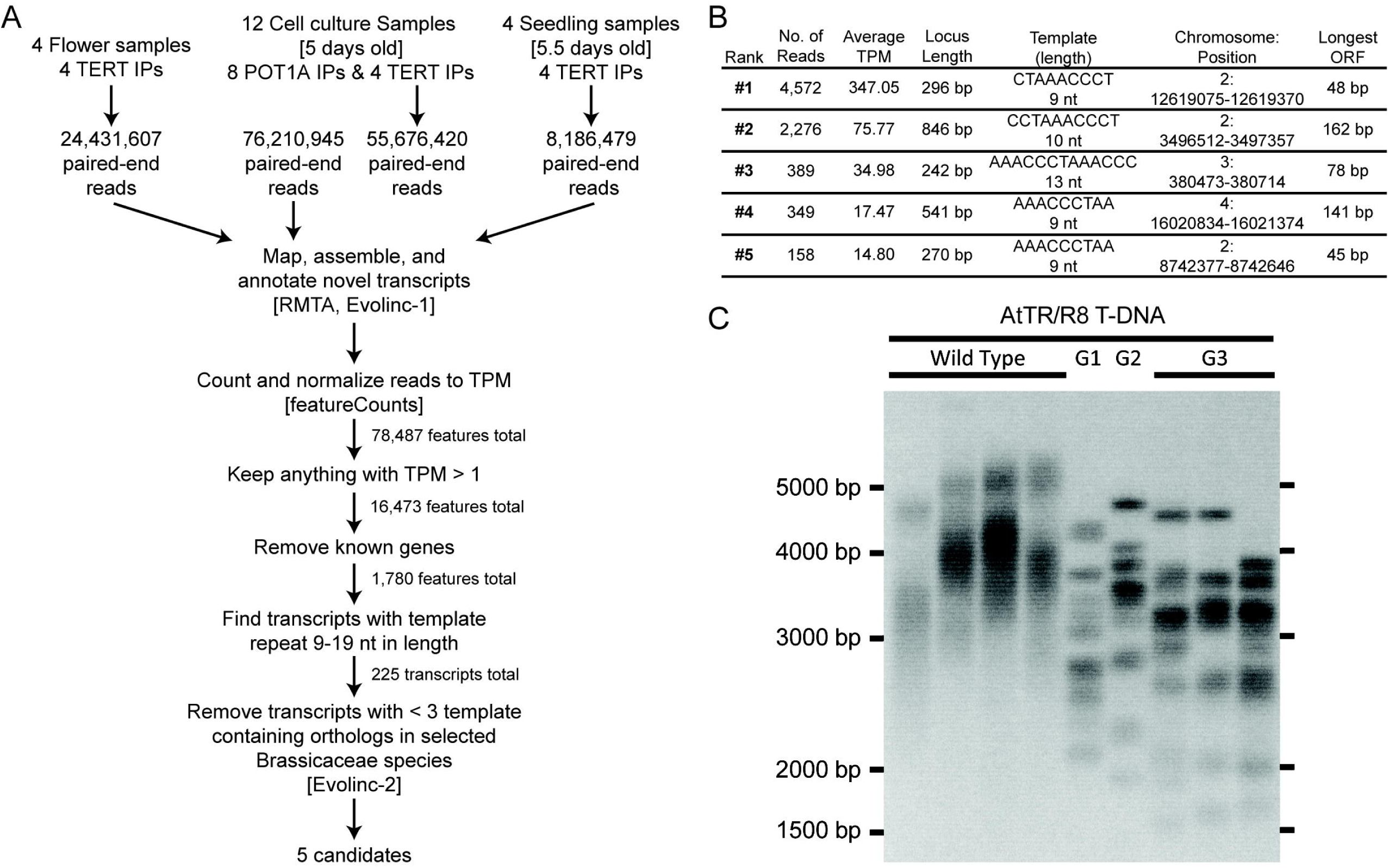
Computational identification of a putative telomerase RNA. (A) Flowchart showing analysis pipeline for RNA-immunoprecipitation. (B) List of the five telomerase RNA candidates identified through the analysis pipeline and ranked by highest average transcripts per million (TPM). (C) TRF of *AtTR/R8* T-DNA insertion lines from wildtype through third generation (G3) homozygous insertion lines with each lane representing separate individuals.

To confirm the involvement of this lncRNA in telomere maintenance, we obtained T-DNA mutants of *AtTR/R8* and confirmed homozygosity. We grew these lines for three generations, measuring telomere length using a TRF with each generation (Fig 5C). We found that homozygous *Attr/r8* mutants showed discrete banding patterns similar to *Attert* mutants. This phenotype is a hallmark of telomerase deficiency [38] and indicates an important role for *AtTR/R8* in telomere maintenance.

### The *AtTER-like* locus is transcribed in other Brassicaceae

The relatively high expression of *AtTER2* and its reported role in the negative regulation of telomerase in *A. thaliana* [15] raises the possibility that the transcript produced from the *AtTER1/2-like* locus may be acting as a telomerase interacting RNA (TIR) similar to *AtTER2*. To address whether the *AtTER1/2-like* loci recovered from other species of Brassicaceae could potentially encode a TIR, we tested whether or not we could detect transcripts consistent with a TIR, but independent of the transcription of the *RAD52-1A* mRNA. For our analyses we chose four species with varying phylogenetic distances from *A. thaliana*: *C. rubella* (diverged ∼20 MYA), *C. hirsuta* (diverged ∼35 MYA), *E. salsugineum* (diverged ∼ 43 MYA), and *S. parvula* (diverged ∼ 43 MYA) [17]. The *AtTER1/2-like* locus partially overlaps *RAD52-1A* in each of these species, therefore we performed RT-PCR on *RAD52-1A* in each species in order to map the intron-exon boundaries (Fig 6A). Following cloning and sequencing of *RAD52-1A* mRNA, we designed reverse primers that bind within the first intron of *RAD52-1A* and used it in combination with a forward primer in the predicted 5’ UTR (Fig. 6A and 6B). We amplified and sequenced transcripts from RT-PCR products generated using this forward primer with a reverse primer in the first intron of *RAD52-1A*. Our results indicate that either a lncRNA is produced at the locus in the tested species or splice variants for *RAD52-1A* exist. Thus, whether the amplified RNAs are entirely distinct from the transcript produced from *RAD52-1A* and if they are TIRs performing a similar role as *AtTER2* remains an open question.

**Fig 6.**
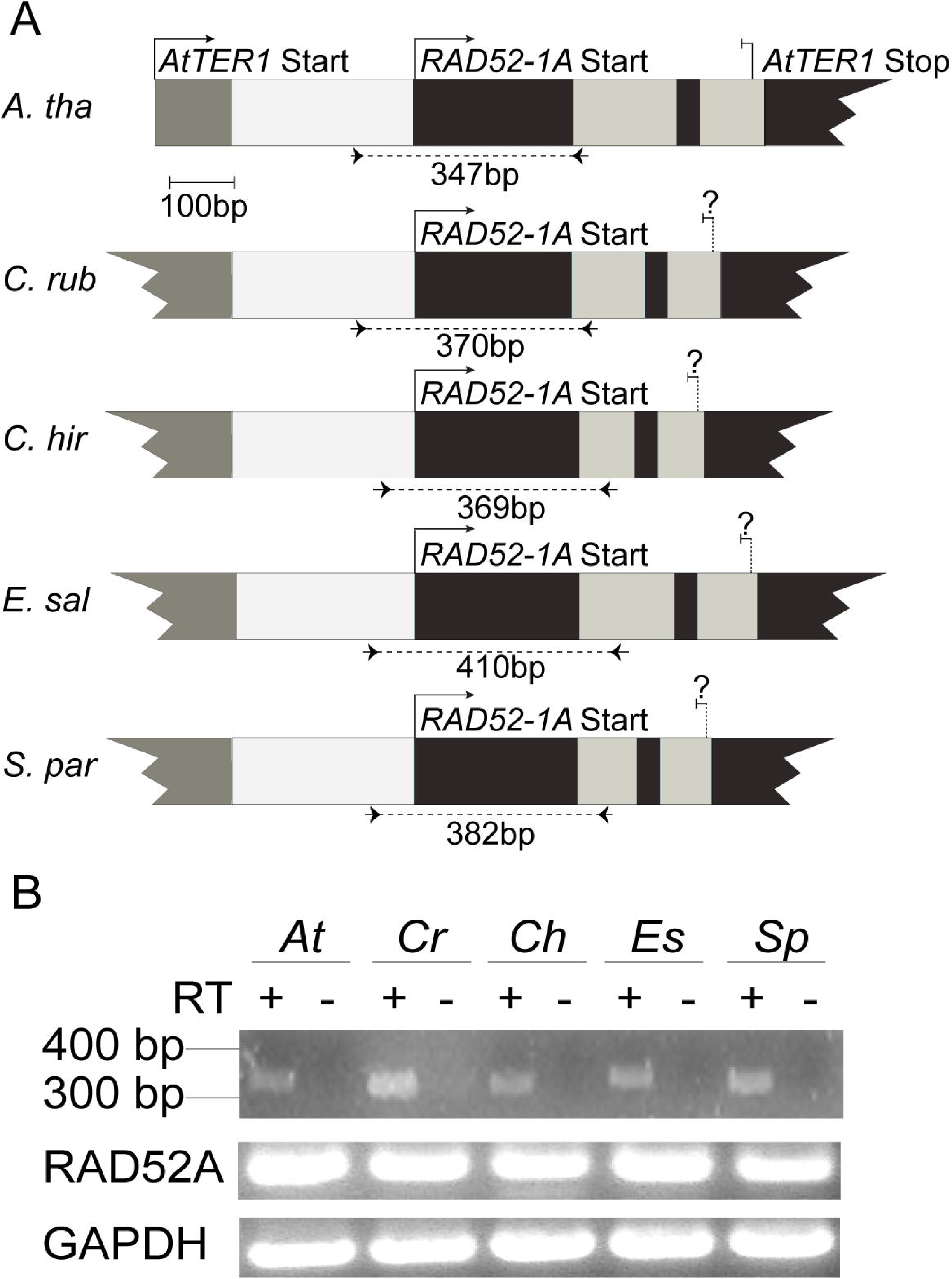
The *AtTER1/2-like* locus is transcribed in four other Brassicaceae species. (A) Schematic of the *AtTER1/2-like* locus in the Brassicaceae species examined in this study. The overlapping *RAD52-1A* locus is also shown. Known transcription start sites (or ones shown in this study) are indicated by a solid arrow. Conserved but unverified TER transcription stop sites are indicated by dashed vertical black lines. Exons are shown by dark grey boxes, while introns are light grey. Primer binding sites for determining putative TER expression are shown as a black arrow beneath each locus. Expected product length is shown. Exon/intron lengths and positions are shown to scale. (B) Transcriptional analysis of the putative TER loci. RT-PCR was performed to determine TER expression using primers designed in the 5’ UTR and within the second intron of *RAD52 1A* as shown in (A). RT-PCR was performed on *RAD52-1A* to determine intron/exon boundaries.

## Conclusions

Our results indicate that the original identification of the telomerase RNA in *A. thaliana* was incorrect, in agreement with the findings of Fajkus et al. (2019). Moreover, our data indicate that the true TER locus is highly conserved in the family, and that the telomerase holoenzyme across the family shares both key protein components as well as biochemical activity. Independently through RNA-seq of AtTERT and AtPOT1A pull downs, we identified *AtTR/R8*, a highly conserved lncRNA within the Brassicaceae that has the potential to act as the canonical TER. These results are further confirmed by recent work in Asparagaceae and a number of other taxa [20]. Fajkus et al. (2019) indicated that the same locus is conserved in a variety of species spanning land plant history, indicating a single evolutionary origin of plant TER. Our work expands on the findings in Fajkus et al. (2019) by showing a direct interaction of this RNA with the telomerase holoenzyme in multiple tissues and stages in development in *A. thaliana*. Further, we showed that despite sequence changes to any potential telomerase RNAs, TERT proteins from across the Brassicaceae family can at least partially rescue an *Attert* mutant. This finding points to the evolutionary stability of the entire telomerase RNP complex. Finally, whether *AtTER2* has a role as a telomerase regulatory RNA remains an open question. We detected transcripts from the *AtTER1/2-like* locus in other Brassicaceae, but were unable to verify if these represent splice variants of *RAD52A-1* or a distinct lncRNA product. One important consideration is that the phenotypes observed for *AtTER2* under genomic stress may be due to a partial protein product encoded from this locus in *A. thaliana*. Regardless, future work on telomerase regulation is required to distinguish between these hypotheses.

## Materials and Methods

### Plant Material and Propagation

*A. thaliana* (Col-0) seed from Dr. Dorothy Shippen, Texas A&M University; *C. rubella* from Dr. Steven Wright, University of Toronto; *C. hirsuta* seed from Dr. Angela Hay, Max Planck Institute for Plant Breeding Research; *E. salsugineum* seed from Dr. Karen Schumaker, University of Arizona; *S. parvula* seed from Drs. Maheshi Dassanayake and Dong-Ha Oh, Louisiana State University were used in this research. Standard Arabidopsis growth conditions were used for all species.

### CRISPR-Cas9 Mutation Analyses

The suite of CRISPR plasmids from Schiml, et al. (2014) were obtained from the Ohio State University Arabidopsis Biological Resources Center (ABRC) and used for cloning of CRISPR-Cas9 and CRISPR-Cas9-D10A vectors, following the procedure described in Schiml, et al. (2014) [44]. A protospacer (5’-GGGTTTAGTTGTCGTCTGAT-3’) overlapping the template domain of *AtTER1* was used in conjunction with both Cas9 and Cas9-D10A; in addition, a second protospacer (5’-TTGTCCGGCGACAGAAATGG-3’) targeting 33 base pairs downstream of the template domain was used with Cas9-D10A. T2’s were screened using PCR of the *TER1* locus followed by HindIII restriction digest for deletions of the entire template domain.

*Atter2* mutants *ter2-2* (SALK_121147) and *ter2-3* (SALK_140126) were obtained from the ABRC then crossed to the two *ter1* alleles, *ter1 Δ22* and *ter1 Δ18*, respectively. The progeny resulting from the crosses were hemizygous for a *ter2* allele and heterozygous for a *ter1* allele. By selfing these individuals we obtained a population segregating the mutations for both genes allowing direct comparisons between full siblings in TRF analyses.

### Nuclear Protein Isolation, TRAP, TRF and RT-PCR

Nuclear extracts were obtained from ∼10g of seedling tissue as described elsewhere [14]. TRAP was performed as described previously [25]. Briefly, 50ng of total protein, sourced from flower tissue, was added to a 25ul final reaction volume containing ∼3uC 32P-dGTP, Gotaq hot start mastermix (Promega), and 400nM Forward TRAP primer. This mixture was placed at 37°C for 45 minutes, followed by the addition of 400nM TRAP Reverse primer. For a list of primers see Supplemental Table 1. PCR followed with an initial 95°C step for 3’, followed by 95°C for 15”, 60°C for 30”, 72°C for 1’ 30” and 35 cycles. TRAP products were resolved on a 6% acrylamide gel (19:1) with 7M urea. For RNA extraction, SDS extraction buffer (final concentration of: 2mM Tris pH 7.5, 200uM EDTA pH 8.0, 0.05% SDS) was added to 1ml of fraction supplemented with enough 3M sodium acetate (300mM final concentration). Following vortexing, this mixture was phenol/chloroform extracted and ethanol precipitated according to standard protocols. Terminal restriction fragment (TRF) length analyses were performed as described in Nelson et al. (2014). In brief, genomic DNA was extracted and digested with the restriction enzyme Tru1I, followed by a Southern blot using the telomeric repeat as a probe. RT-PCR was performed as described above using primers listed in Supplemental Table 1.

### Positive Selection Analyses

We obtained CDS sequences for *TERT, Dyskerin, Ku70*, and *Ku80* from 15 taxa (Brassicaceae or close relatives) from publicly available genomes using CoGe, TAIR, Phytozome, and SALK [41,45,46]. *Branch-sites* likelihood ratio tests were performed on the indicated branches of each gene tree using PAML (codeml) [24] to compare the likelihood scores of models of evolution that either include site classes with ω values ≥ 1 (denoted Alt = Alternative model) or explicitly exclude these site classes (denoted Null = null model). Alternative model Likelihood scores that are ≥ 1.92 better than null model Likelihood scores indicate a significant signal of positive selection (p < 0.05) along the specified branch (see critical values from chi-squared distribution table at one degree of freedom). For *Ku70* and *Dyskerin, L. alabamica* was removed from our analysis because the inferred phylogeny was not congruent with the accepted organismal phylogeny. The *C. hirsuta* branch was not tested for *Ku70*/*80* and *Dyskerin*, as genomic sequence was not available for these genes.

### Substrate Specificity Analysis

Substrate specificity was tested by replacing the TRAP Forward G primer with alternative forward primers, keeping the reverse primer the same. These products were resolved on a 6% acrylamide sequencing gel. The space between two bands generated from the N15-GGG oligo was divided into seven quadrants (using ImageJ, NIH). Where bands in other lanes migrate relative to these quadrants determines the value given to them for “observed product difference.”

### *tert* -/- Complementation

Full length genomic *TERT* constructs were PCR amplified from each species tested. These constructs were designed to include 2 Kb upstream and 0.5 Kb downstream sequence to include appropriate regulatory regions. Constructs were cloned into the promoter-less Gateway (Invitrogen) vector pH7WG using In-Fusion (Clontech) and then sequence verified. *Agrobacterium tumefaciens* (strain GV3101) was used to transform this binary vector into a fifth generation *tert* -/- *A. thaliana* background using the floral dip method [47]. At least three independent transformed lines were identified for each construct. RT-PCR was performed on each line to ensure transcription of the *TERT* transgene using primers denoted in Supplemental Table 1. PETRA was performed as described in Heacock et al. (2004) using subtelomeric primers specific for chromosome arms 1L, 3L, and 5L (see S1 Table for primer sequence). Radioactive images were quantified using ImageJ (NIH).

### RNA Immunoprecipitation (RIP) of the Telomerase Complex and RNA-seq Library Preparation

5.5-day old *A. thaliana* accession Columbia (Col-0) seedlings, 5-day old *A. thaliana* T87 cell culture, and *A. thaliana* Col-0 flowers were used for IP experiments. Cell culture was concentrated using Miracloth and the resulting powder was dried, packed, and stored at −80 °C for IP. Samples were pulverized in liquid nitrogen and approximately 100 mg of tissue powder was resuspended in 5 mL of cold RIP buffer (200 mM Tris-HCl pH 9, 110 mM potassium acetate, 0.5 % Triton X-100, 0.1 % Tween 20, 2.5 mM dithiothreitol (DTT), 1.5 % Protease Inhibitor Cocktail (Sigma-Aldrich P9599), 40 units/μL RNaseOUT (ThermoFisher p10777019)) by pipetting. The 5 mL of homogenate was then centrifuged at 1,500 × g for 2 minutes at 4°C. From the resulting supernatant, 250 μL was kept as RIP input, and 5 μL was kept for the TRAP assay.

Immunoprecipitation was performed by washing 150 μL of Protein A Dynabeads (ThermoFisher, 10002D) twice with PBS (137 mM NaCl, 2.7 mM KCl, 10 mM sodium phosphate dibasic, 2 mM sodium phosphate monobasic, pH 7.4) supplemented with 0.02% Tween 20. Then, 5-10 μL of GFP (Abcam, ab290), TERT, or POT1A antibody was diluted to 600 μL with PBS supplemented with 0.02 % *Tween* 20, 0.2 mg/mL Salmon Sperm DNA (ThermoFisher, 15632011), and 0.25 mg/mL BSA. Washed beads were resuspended in the antibody solution and allowed to rotate at room temperature for 90 minutes.

After 90 minutes, the beads were separated from the antibody-PBS solution and resuspended in 2 mL of the RIP extract. The bead-RIP mixture was rotated at 4°C for 3 hours. After 3 hours, beads were washed 6 times with 1 mL cold RIP buffer and washed twice with 1 mL of cold TMG buffer (10 mM Tris-Acetate pH8, 1 mM MgCl_2_, 1 mM DTT, 10 % glycerol). Beads were resuspended in 150 μL of cold TMG buffer, 5 μL was kept for TRAP assay.

To purify the RNA associated with the telomerase RNA complex, we added 1 mL of TRIzol Reagent (ThermoFisher, 15596026) to the 145 μL of remaining beads at the end of the RIP experiment to extract RNA following manufacturer’s instructions. RNA was treated with TURBO DNase (ThermoFisher, AM2238) to degrade contaminating genomic DNA following the manufacturer’s instructions. RNA was purified from the DNase reaction by ethanol precipitation. The RNA libraries were prepared using the YourSeq FT v1.5 kit (Amaryllis Nucleics) following manufacturer’s instructions. RNA samples checked for integrity using a Bioanalyzer throughout preparation. RNA was sequenced using Illumina NextSeq 500 PE × 75bp.

### Computational Analysis of RNA-seq Data

RNA purifications from POT1A and TERT antibodies generated 164,505,451 total paired-end reads across 20 pull-down experiments. Reads were mapped to the *A. thaliana* TAIR 10 reference genome using the Read Mapping and Transcript Assembly workflow (RMTA v2.5.1.2) (github.com/Evolinc/RMTA) through the CyVerse Discovery Environment (cyverse.org). RMTA utilizes Hisat2 [48] for read mapping, and Stringtie [49] for transcript assembly as a seamless workflow. Default RMTA parameters were used, besides specifying paired-end reads and changing the maximum intron length from 500,000 bp to 50,000. Read mapping for all experiments achieved at least 89% overall alignment rates. Candidate loci were identified using Evolinc v1.7.5 [42]. Evolinc identifies and annotates novel long intergenic noncoding RNAs (lincRNAs) using assembled transcripts from RMTA as GTF files as well as information regarding surrounding annotated genes using a reference genome annotation. Candidate lincRNAs from the 20 pull-down experiments were merged with the reference genome annotation to a single annotation file using Evolinc merge for subsequent analyses.

To quantify the expression of candidate loci, featureCounts v1.6.0 [50] was used (specifying paired-end reads, feature types: exon, and gene attribute: gene_id) taking BAM files from RMTA as input along with the merged annotation file. Gene expression was normalized across the 20 pull-down experiments by converting counts to transcripts per million (TPM) which accounts for feature length and sequencing depth.

To begin filtering for candidate telomerase RNAs, genes were only selected if they had an average TPM (Transcripts per Kilobase Million) over 1 across the 20 experiments (78,487 features to 16,473 features). Candidates overlapping known genes, from TAIR 10, were then discarded (16,473 features to 1,780 features). Candidate loci were then analyzed for putative template domains by mapping all permutations of the telomere repeat “TTTAGGG” and its reverse complement from 9 to 19 nt in length (154 possible permutations) to the 1,780 candidates using the software Geneious Prime 2019.2.1 (BioMatters, Inc.), resulting in 225 remaining candidate loci containing at least a 9 nt template domain.

Conservation analyses on the 225 candidate loci was performed using Evolinc-II which identifies orthologous loci across a defined set of genomes, in this case from representative species across the Brassicaceae family *(Arabidopsis thaliana, Arabidopsis lyrata, Capsella rubella, Brassica oleracea, Brassica rapa, Eutrema salsugineum, Schrenkiella parvula*, and *Aethionema arabicum*) as well as one species from the sister family, *Tarenaya hassleriana* in Cleomaceae. Evolinc-II was used with an E-value cutoff of E-5 to allow for sequence divergent orthologs to be identified. Finally, TER candidates were removed if a template containing ortholog was not identified in at least 3 relatives, yielding 5 candidates.

### Telomere Length of AtTR/R8 T-DNA Lines

The T-DNA line (FLAG-410H04) was obtained from the Versailles INRA collection. Plants homozygous for the T-DNA were bred for three generations to show the long term impact of a *AtTR/R8* loss on telomeres. The wildtype comparison used for the TRF was a homozygous wildtype sibling obtained from a population segregating for the FLAG-410H04 T-DNA insertion.

## Acknowledgements

We thank Dr. Dorothy Shippen for the use of anti-TERT and anti-POT1A antibodies. The efforts we describe to identify and characterize the true telomerase RNA in Arabidopsis benefitted from collaboration with the Shippen Lab at Texas A&M University. In particular we thank Jiarui Song and Claudia Castillo Gonzalez for sharing data and results related to the identification and characterization of the Arabidopsis TR. They’re findings are described in an independent manuscript (Song et al.) currently under revision. Finally, we thank the PaBeBaWoMo research group and members of the Beilstein Lab at the University of Arizona for insightful and helpful comments during the completion of this project.

## Figure Legends

**Supplemental Fig 1. Positive selection analysis of Dyskerin, Ku70, and Ku80 in Brassicaceae**. Roman numerals correspond to branches tested in Fig 2. Alternative model likelihood scores that are significantly greater than null model likelihood scores indicate positive selection (p<0.05) along the specified branch and are denoted in bold.

**Supplemental Fig 2. Substrate utilization of telomerase from different Brassicaceae telomerase RNPs**. (*A*). Expected binding between AtTER1 template and each oligo used in the experiment. The first set of nucleotides added by this template before the first full repeat is shown in bold. Expected product differences are calculated relative to oligo #4, which is the shortest oligo provided in the assay and therefore serves as the baseline. (*B*) A subset of the gel image from the substrate utilization assay for each species tested. A dashed line is drawn through the center of a band for oligo N15-GGG, which serves as the baseline for calculating observed product differenced for the other oligos.

**Supplemental Fig 3. Test of complementation of *Attert* -/- background with genomic TERT constructs from other Brassicaceae**. (*A*) PETRA results for Attert -/- lines transformed with *C. rubella* TERT (Cr), *C. hirsuta* TERT (Ch), and *E. salsugineum* TERT (Es). (*B*) Quantification of results shown in (*A*).

**Supplemental Fig 4. Confirmation of expression of TERT transgenes for complementation experiments**. RNA was extracted from floral tissue from each background. RT-PCR was performed on both the 3’ end of the TERT transgene (shown) and the full-length construct (not shown). GAPDH was used to determine quality of RNA and as an approximate loading control. “+” indicates a positive RT reaction, whereas “-” indicates no reverse transcriptase was added.

## Notes

#### Summary of Updates

This revision updates the main body and acknowledgements to highlight the contributions of members of the Shippen Lab at Texas A&M University on the results reported.

