## Supplemental Figures for "Evolutionary and biochemical analyses reveal conservation of the Brassicaceae telomerase ribonucleoprotein complex"

Supplemental Figure 1

| Gene | Branch | Null (ω ≤ 1) | Alt (1 ≤ ω) | Alt-Null | Liklihood ratio score (2[Alt-Null]) | Significance |
|---|---|---|---|---|---|---|
| Dyskerin | i | -6102.87 | -6102.87 | 0 | 0 | p > 0.05 |
| | ii | -6102.85 | -6102.85 | 0 | 0 | p > 0.05 |
| | iii | -6100.38 | -6096.32 | 4.06 | **8.13** | **p ≤ 0.01** |
| | iv | ND | ND | ND | ND | ND |
| | v | ND | ND | ND | ND | ND |
| | vi | -6096.37 | -6094.80 | 1.57 | **3.14** | **p ≤ 0.05** |
| Ku70 | i | -5799.49 | -5799.49 | 0.00 | 0.00 | p > 0.05 |
| | ii | -5798.94 | -5796.68 | 2.26 | **4.52** | **p ≤ 0.05** |
| | iii | -5799.49 | -5799.49 | 0.00 | 0.00 | p > 0.05 |
| | iv | ND | ND | ND | ND | ND |
| | v | ND | ND | ND | ND | ND |
| | vi | -5799.49 | -5799.49 | 0.00 | 0.00 | p > 0.05 |
| Ku80 | i | -7725.67 | -7725.67 | 0.00 | 0.00 | p > 0.05 |
| | ii | -7725.67 | -7725.67 | 0.00 | 0.00 | p > 0.05 |
| | iii | -7725.67 | -7725.67 | 0.00 | 0.00 | p > 0.05 |
| | iv | -7724.92 | -7724.92 | 0.00 | 0.00 | p > 0.05 |
| | v | ND | ND | ND | ND | ND |
| | vi | -7725.67 | -7725.67 | 0.00 | 0.00 | p > 0.05 |

Supplemental Figure 2

**A**

| | Expected Product Difference (EPD) Relative to N15-GGG | | Observed (for *A. thaliana*) |
|---|---|---|---|
| 1: | N15-NNNNAG**GG**<br>3′-AUCCCAAAUC-5′ | **= +5** | **+4** |
| 2: | N15-NNNNGG**G**<br>3′-AUCCCAAAUC-5′ | **= +4** | **+4** |
| 3: | N15-NNNGGG<br>3′-AUCCCAAAUC-5′ | **= +3** | **+3** |
| 4: | N15-GGG<br>3′-AUCCCAAAUC-5′ | **= +0** | **+0** |
| 5: | N15-NNNNTT**TAGGG**<br>3′-AUCCCAAAUC-5′ | **= +1** | **+1** |
| 6: | N15-NNNNNT**TTAGGG**<br>3′-AUCCCAAAUC-5′ | **= +2** | **+6** |
| 7: | N15-NNT**TTAGGG**<br>3′-AUCCCAAAUC-5′ | **= +6** | **+3** |

**B**

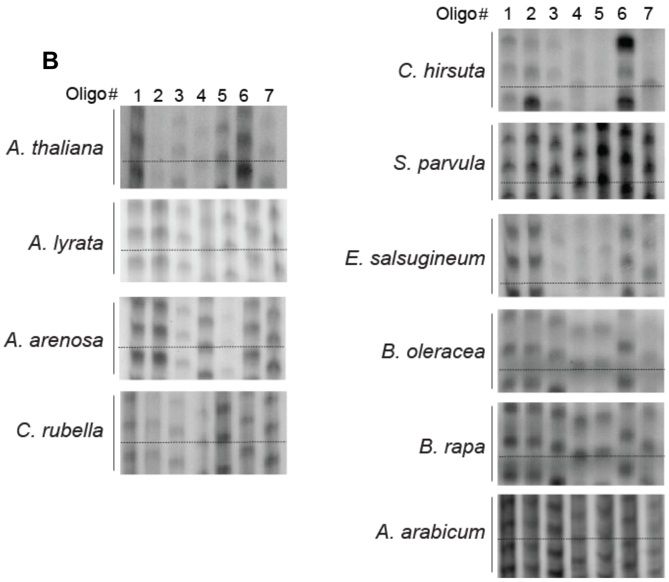

Supplemental Figure 3

A

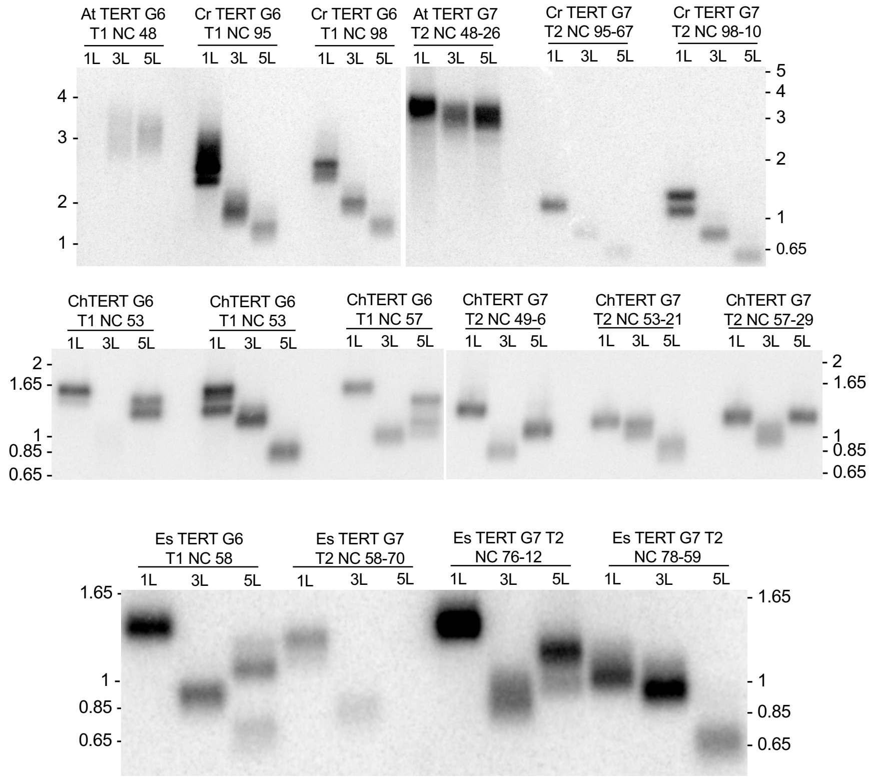


Supplemental Figure 3

B

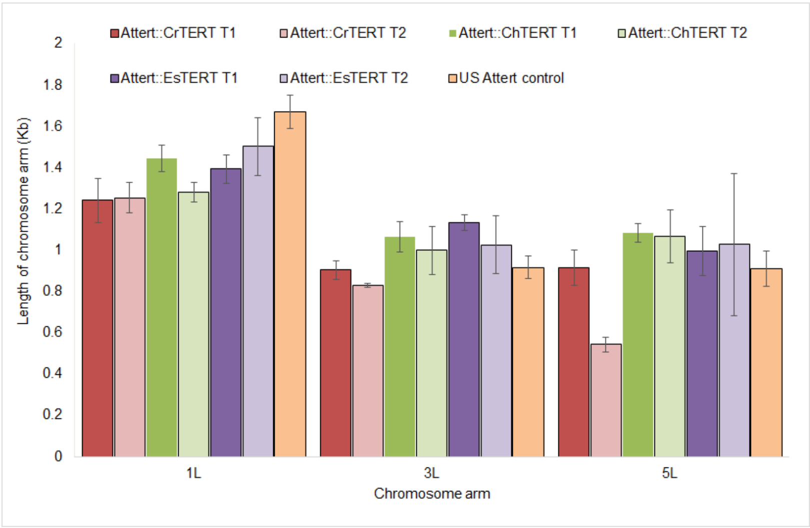

Supplemental Figure 4

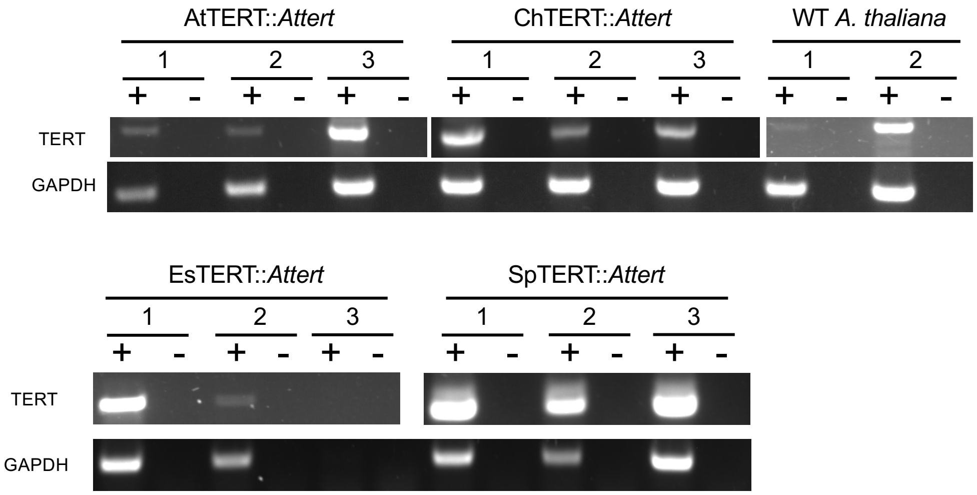
